# Multispecies plastic associated bacterial community MAGs separation by electromagnetic radiations

**DOI:** 10.1101/2025.08.21.671670

**Authors:** Harshal S. Jadhav, Abhay B. Fulke, Chandrashekar Mootapally, Neelam M. Nathani

## Abstract

Plastic is a pollutant that is believed to be difficult to remove from the environment. Most portion of the plastic is a hydrocarbon, which can act as a substrate for the attachment of microorganisms and serves as a source of nutrition for attached microorganisms. We observed and investigated microbial communities developed on *in situ* incubated low-density polyethylene for 6 months in the ambient marine environment. The plastic degrading microbial community formed was given electromagnetic radiation pretreatment to enhance the recovery of microbial community involved in plastic degradation using shotgun metagenomic sequencing and the probable mechanism of separation of plastic degrading microbial community involving the presence of photoreceptor which sense particular wavelength electromagnetic radiation and activate gene inhibitor. Four different phyla such as Actinobacteria, Bacteroidetes, Firmicutes, and Proteobacteria have been examined from a total of 62 metagenome-assembled genomes (MAGs). Phylum Proteobacteria was observed to be abundant plastic degrading bacteria; whereas other phyla might be symbiotically associated with proteobacteria in plastic combinely form the plastic associated bacterial community. This is a first report that deals with the community pattern of multispecies plastic degrading bacterial community separated from a substrate by electromagnetic radiations of 650 nm (red light) and 480nm (blue light). Bacterium *Piscirickettsiaceae* specifically and abundantly (46.57) response to red light whereas bacterium *Chromatiaceae* specifically and abundantly (16.99) response to blue light.

**Importance:** In natural marine environment especially at benthic region, microbes can efficiently degrade plastic by making biofilm on its surface. However this strongly attached biofilm is very much difficult to detach from its substratum. Some people even used high pressure water or scrap using sharp surface in order to remove this biofilm from plastic. However major drawback of using such techniques is we can not recover this biofilm in intact form. In present study we used application of different wavelength electromagnetic radiations in order to recover this biofilm in intact form. It is based on the observation that the bacterial density significantly decreases at higher intensity of light.

**Synopsis:** This study significantly demonstrate strongly attached (on the surface of plastics) plastic degrading marine microbial community composition and function on pretreatment with electromagnetic radiations.

## Introduction

More than 380 million tons of plastic is being produced globally every year and around 8 million tons, which is 3% of total plastic waste generated each year enters the oceans (Jambeck et al., 2015). Due to physical, chemical and biological factors, plastic enter the ocean converted into microplastics, which are even more difficult to remediate (Narwal et al., 2024). The consequences of plastic in the ocean ecosystem are not well known. Other than this, around 55% of total plastic waste is discarded in the environment as such, 20% of plastic waste is recycled and 25% of plastic waste is incinerated (Geyer et al., 2017). This waste plastic discarded in the ocean ultimately meet the ocean via river water or sewage or rain water run off. Hence studying marine based bacteria that can degrade plastics with higher efficiency would be the essential task to develop potential plastic remediation strategies.

A large amount of plastic enters the oceans from land, vessels and beachgoers (Pruter 1987). Microorganisms in the marine environment have adapted plastic as a source of nutrients. These plastic hydrocarbons are extremely slow to degrade (Zettler et al., 2013). Marine plastic biofouling is an attachment of biotic and abiotic factors on the surface of plastic so that it sinks downward in the water column (Urbanek et al., 2018). Bacterial community form on the plastic surfaces immediately once plastic enter the marine environment or water body (Bhagwat et al., 2021), providing a platform for the attachment of microalgae, microphytobenthos and other microorganisms, further forming biofouling communities. These plastic degrading bacteria mineralizes plastic to carbon dioxide (Rummel et al., 2017; Jadhav et al., 2024). Plastic hydrophobicity also decreases as it sinks down due to the attachment of biofouling community (Lobelle and Cunliffe 2011). The rate of plastic degradation is found to be higher in the benthic zone compared to the planktonic zone of marine environment (Abed et al., 2020).

Regulation of plastic degrading bacterial community (which is firmly attached to its substratum) is a problematic task. Even detachment of this bacterial community requires harsh treatment (Kirstein et al., 2019). Due to this, we may not recover bacterial community in intact form. In present study, different wavelength electromagnetic radiation pretreatment is given to in situ incubated plastic to enhance recovery of plastic degrading bacterial community specific to particular wavelength. To sense light (photons) of particular wavelength, there must be involvement of photoreceptor. This photoreceptor activates DNA repressor, which stops synthesis of proteins involved in bacterial attachment to plastic. Since plastic is a compact molecule which mainly prepared from petroleum hydrocarbons. These plastic hydrocarbons can serve as a huge source of nutrients to microorganisms in biofilm. Hence excess of these nutrients must be store inside the microbial cells in the form of lipids.

Recently, Jadhav et al., 2024 showed that plastic degrading bacterial group can transition from a collective community to free-living state in response to light. In context to the same, we designed a light exposure pretreatment setup to separate the bacteria containing similar photosensitive machinery, so that diverse bacteria can be found on plastic surface which can be characterized at a higher resolution. The objectives of this study were to determine plastic degrading microbial community composition and their function, pretreatment with electromagnetic radiation; to predict a collective model for bacterial attachment over plastic in situ condition in the marine environment derived by analyzing plastic-associated MAGs and functional potential using shotgun metagenomics.

## Material and Methods

### *In situ* incubation of polyethylene

Approximately 0.5 mm thickness low-density polyethylene or LDPE was fitted in three different stainless-steel frames of 15cm × 15cm separately and incubated underwater at sea surface at latitude- 19.49048 and longitude- 72.82067 as shown in figure 1. Samples were incubated at bottom or sea surface for six months from September 2020 to February 2021. The selected location is such that the water at the location never dries (Pinto et al., 2019). Before incubation, LDPE films were rinsed with 70% isopropanol and Milli-Q water. LDPE and water samples were brought to the laboratory in a proper aseptic condition and immediately processed.

**Figure 1.**
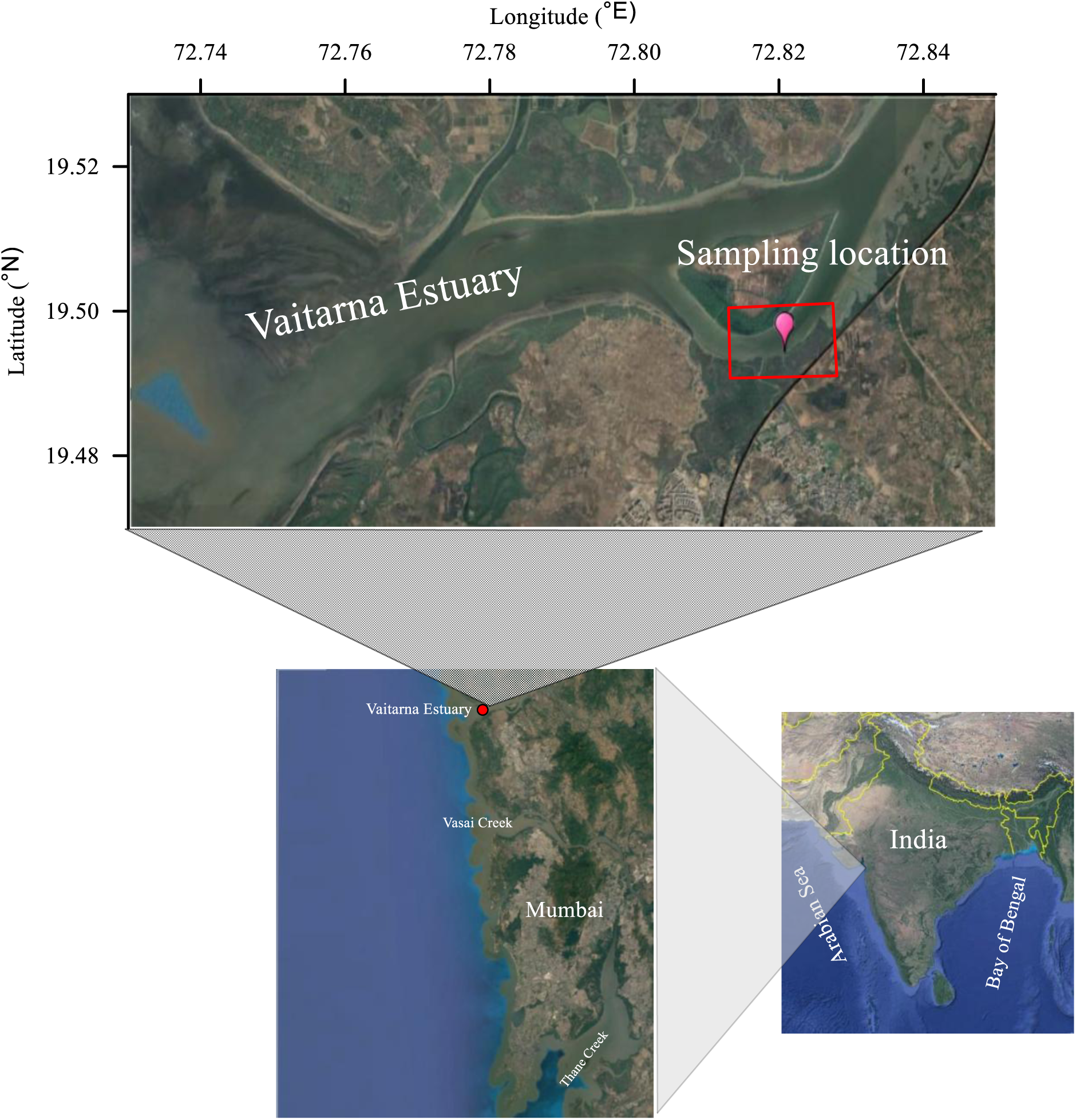
Sampling location of *in situ* incubated polyethylene

### Physicochemical parameters analysis

Physicochemical parameters of the seawater collected in February 2021 from the same plastic incubation site were measured using the standard protocol as described in Fulke et al. 2019.

### Light intensity and wavelength measurements

Light intensity was measured using Lutron LX-101A digital lux meter. Values obtained were converted to µmolem^-2^s^-1^ using conversion of 1Klux= 19.5 µmolem^-2^s^-1^ (Fulke et al., 2015). Light wavelength was measured using spectrofluorometer (RF-6000: SHIMADZU).

### Electromagnetic radiation pretreatment to assess bacterial diversity

After *In situ* incubation, LDPE film developed with natural plastic associated bacteria was cut into 4cm × 4cm strips aseptically. Four such plastic strips per flask were inoculated in three different 1000 ml Erlenmeyer flasks along with 400 ml autoclaved 0.8% saline solution in each flask. These flasks with plastic strips and saline were then incubated for 48 hours in three different wooden chambers (base: 15 cm; height: 40 cm) separately under room temperature around 23-28^0^C. Among three such chambers, one was kept dark or controlled (C), and the other two PE-1 and PE-2 were fitted with Light Emitting Diode (LED) for different electromagnetic radiation intensity around 650 nm for red and 480nm for blue, respectively. The constant intensity of around 230 µmole m^-2^ s^-1^ with 4-watt LED lights were maintained in the PE-1 and PE-2 chambers. During the incubation, the light intensity and wavelength of LED was monitored using Lutron LX-101A digital lux meter and spectrofluorometer RF-6000: SHIMADZU, Japan, respectively.

### DNA extraction and sequencing

After resuspension of plastic associated bacteria for 48 hours, 150 ml from three biological replicates (C, PE-1 and PE-2) (Blainey et al., 2014) flasks was filtered using Millipore 0.22 µm filter papers from the C, PE-1 and PE-2. Under proper aseptic conditions, the filter paper was cut into small pieces using sterile scissors followed by total community DNA extraction from resuspended plastic associated bacteria of each flask using Qiagen DNeasy powersoil pro kit as per the manufacturer’s protocol. Upon confirmation of quantity and quality using Nanodrop2000, the extracted DNA samples were sequenced by for shotgun metagenomics at MedGenome Labs Ltd., Bengaluru, with a chemistry of 150 x 2 bp using Hiseq4000 Illumina platform. Samples C-3, PE-1 and PE-2 were sequenced using HiseqX with a read length of 151 bp. The V3-V4 hypervariable regions of the DNA was amplified using reverse and forward primers. Further, the samples were processed for whole genome metagenome analysis.

### Quality filtering and metagenome assembly generation

Trim galore (Marcel 2011) was run with default parameters on raw reads to remove adapters and low quality reads. Best Match Tagger or BMTagger (Rotmistrovsky and Agarwala 2011) was used with human genome GRCh38 to remove human contamination from the reads. Final QC passed reads were taken to generate individual sample metagenomic assemblies and a pooled assembly using MEGAHIT (Dinghua et al., 2015) with meta-sensitive parameter that uses kmer list: 21, 29, 39, 49, 59, 69, 79, 89, 99, 109, 119, 129, and 141.

### Genome binning and bin refinement

Metagenome Assembled Genomes (MAGs) were binned using MaxBin2 (Yu-Wei et al., 2016) and CONCOCT (Alneberg et al., 2014) from “C or without light treatment detached plastic associated bacteria, PE-1 or detached plastic associated bacteria by red light treatment and PE-2 or detached plastic associated bacteria by blue light treatment” metagenomic sequence reads individually by considering pooled assembly as reference. Further, individual assemblies of C, PE-1 and PE-2 were used for binning with metaBAT2 (Kang et al., 2019). MAGs of respective sample were consolidated using metaWRAP’s bin refinement module (Uritskiy et al., 2018). The MAGs were subjected for quality assessment using checkM (Parks et al., 2015) and MAGs with <50% completeness and >10% contamination were not considered. Resulting MAGs with >90% completeness and <5% contamination were considered as high in quality, those with >75% completeness and <10% contamination were considered as medium in quality, and rest of the MAGs were considered as low in quality (Johansen et al., 2022).

### Statistical analysis

Statistical analysis was performed using STAMP (Parks et al., 2014). Each of the sample i.e., C, PE-1 and PE-2 were compare among themselves using STAMP software for statistical interpretation.

### Identification, and functional annotations

Individual MAGs were identified by running megablast on NCBI’s Nucleotide database and functionally annotated using DRAM pipeline (Shaffer et al., 2020).

## Data Availability

Raw data is submitted to European Nucleotide Archive (ENA) under the study number PRJEB48191. MAGs are submitted to NCBI under bioproject ID PRJNA773851 and biosample IDs are mentioned in table 1.

**Table 1:** Biosample IDs for 62 MAGs submitted at NCBI.

| Sample | Biosample IDs | Numbers of MAGs |
| --- | --- | --- |
| C | SAMN23076163 to<br>SAMN23076185 | 23 |
| PE-1 | SAMN23076186 to<br>SAMN23076205 | 20 |
| PE-2 | SAMN23076206 to<br>SAMN23076224 | 19 |

## Results and Discussions

Total 62 Bacterial MAGs were extracted from the samples, which included 23 from C, 20 from PE-1, and 19 from PE-2. These MAGs span over four different phyla-Actinobacteria, Bacteroidetes, Firmicutes, and Proteobacteria. Phylogenetic tree is as shown in figure 2a signifies about the total of 62 MAGs from overall light treated and non-treated plastic degrading bacterial community. One of the unclassified gammaproteobacteria have found to be present in both treated samples however absent in without light treated bacterial community (Figure 2b). *Chromatiaceae*, also a proteobacteria was observed to specifically and abundantly in blue light treated sample. Other species like *Azoarcus pumilus*, *Rhodobacteraceae* and *Pseudomonas* were observed to be responsive specifically to blue light while *Thauera* species, *Alteromonadaceae* and *Zoogloeaceae* responded specifically to red light. Therefore, unclassified gammaproteobacteria, *Chromatiaceae, Azoarcus pumilus*, *Rhodobacteraceae*, *Pseudomonas, Thauera* species, *Alteromonadaceae* and *Zoogloeaceae* are predicted to be potential candidates involved in plastic biodegradation and plastic degrading bacterial community formation.

**Figure 2.**
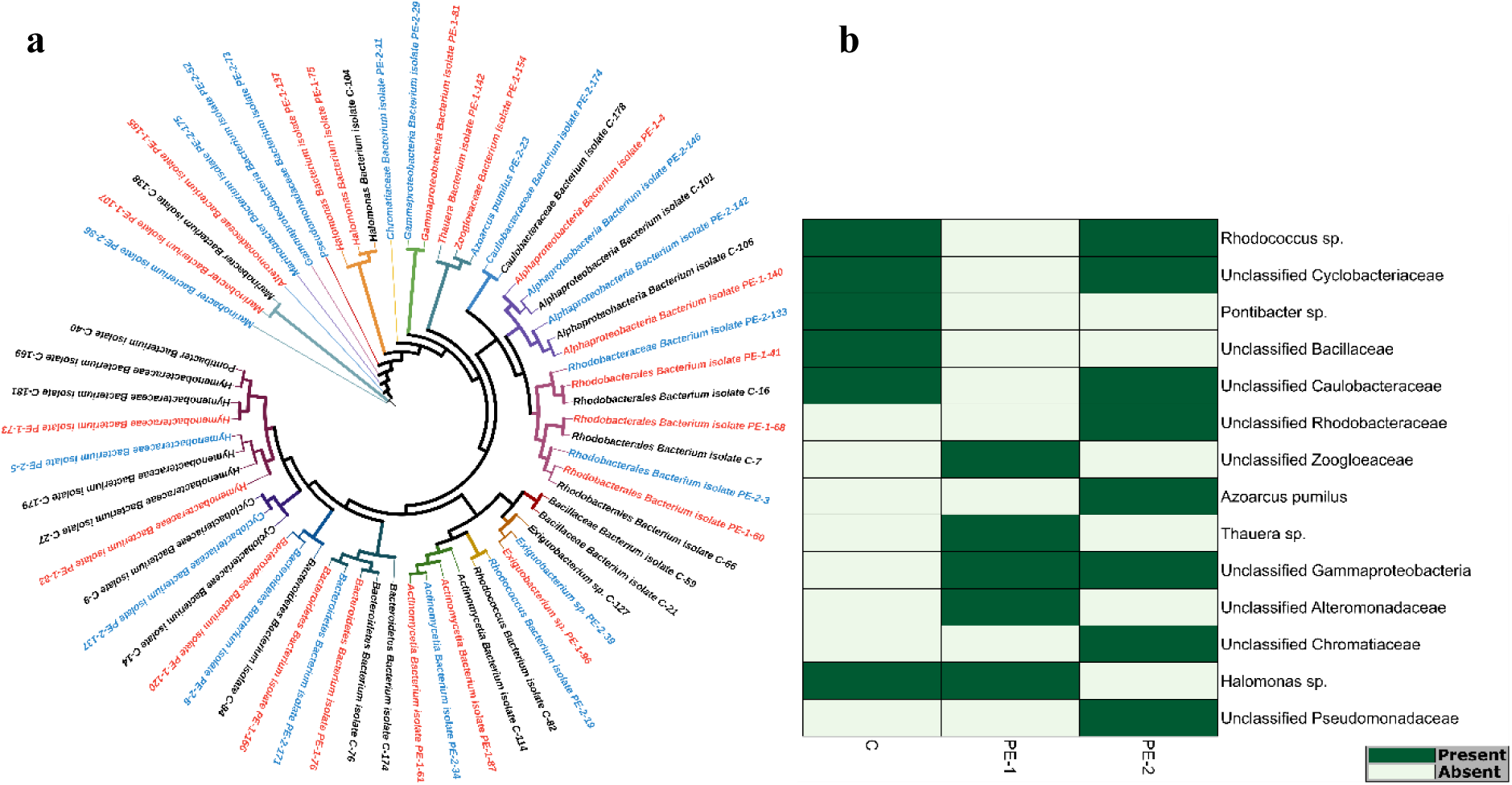
**a** Phylogenetic tree of plastic biofilm associated microbial community MAGs. Red highlighted organisms are red light treated bacterial community, blue highlighted organisms are blue light treated bacterial community and remaining are control or without light treatment for plastic associated bacterial community. **b** Heatmap showing differential presence between biological replicates whereas core microbes present in all three studied samples were excluded from the heatmap.

### Prevailing marine environment of in situ incubation site

As per calculated physicochemical parameters of water where natural microcosm experiment or in situ incubation of plastic was carried out, the water was saline and slightly alkaline hence bacteria involved in plastic biodegradation in this case must be belonging to alkalihalobacteria lineage (Table 2). A large number bacteria from the plastic seems to involve in denitrification hence concentration of nitrite and nitrate are less as compared to ammonia. These processes might symbiotically support plastic biodegradation. Collected water was turbid, and also contain suspended solids or particulate organic matter hence bacteria able to attach on solid surface like plastic must be present in abundant. Polyhydrocarbon or PHc concentration observed is significant and plastic is also a type of polyhydrocarbon therefore presence of PHc degrading bacteria is expected.

**Table 2:** Water quality parameters from the site of plastic incubation.

| Parameter* | SR-1 (A) | SR-1 (B) |
| --- | --- | --- |
| pH | 7.91 | 7.88 |
| Suspended Solids (mg/l) | 47.14 | 51.86 |
| Turbidity (NTU) | 23.50 | 27.10 |
| Salinity (PSU) | 32.48 | 32.53 |
| DO (mg/l) | 6.88 | 6.64 |
| BOD (mg/l) | 3.10 | 2.63 |
| PO<sub>4</sub><sup>3-</sup>-P (μmol/l) | 3.21 | 3.87 |
| NO<sub>3</sub><sup>-</sup>-N (μmol/l) | 10.31 | 10.63 |
| NO<sub>2</sub><sup>-</sup>-N (μmol/l) | 2.43 | 2.77 |
| NH<sub>4</sub><sup>+</sup>-N (μmol/l) | 17.10 | 18.12 |
| Si (OH)<sub>4</sub> (μmol/l) | 10.95 | 12.64 |
| PHc (μg/l) | 15.01 | 13.96 |
| Phenol (μg/l) | 34.65 | 30.80 |

\* Samples collected in duplicate viz. SR-1 (A) and SR-1 (B)

As given in figure 3, species belonging to *Actinobacteria* phylum observed in our data can be divided into two groups: Unclassified *Actinobacteria* and *Rhodococcus* sp. It is evident that *Rhodococcus* sp. can use wide range of metabolic processes than Unclassified *Actinobacteria*. It is also clear that *Rhodococcus* sp. harbors diverse set of carbohydrate active enzymes compared to other Actinobacteria in this bacterial community. Another interesting observation is that nitrogen metabolism is present in all *Actinobacteria* but unclassified *Actinobacteria* converts nitrite to ammonia whereas *Rhodococcus* sp. converts nitrite to nitric oxide and these processes might support actual plastic degradation or helpful in establishment of optimum environment for plastic biodegradation. It is also found that *Rhodococcus* sp. have the arsenate reduction machineries. All actinobacteria are also having more than one SCFA and alcohol conversion reaction.

**Figure 3.**
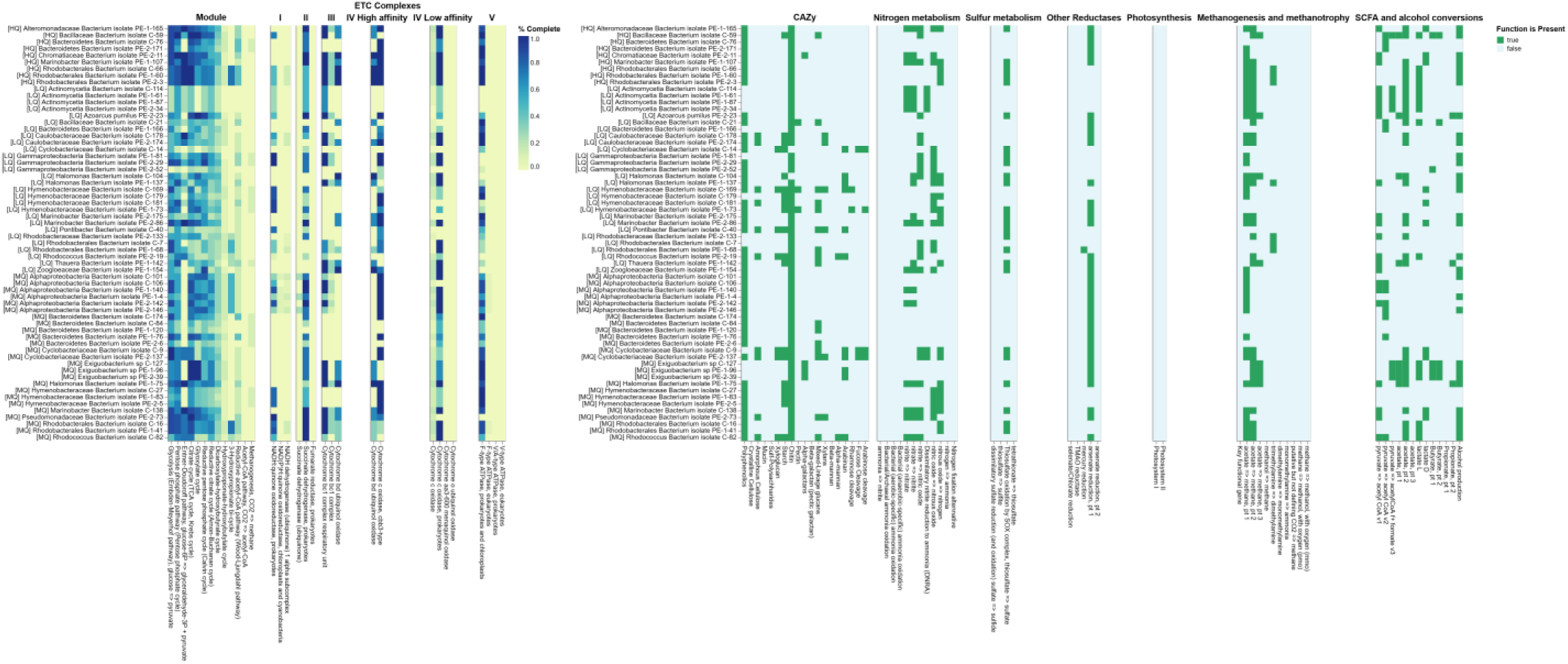
Functional association among species in four different phyla- *Actinobacteria*, *Bacteroidetes*, *Firmicutes* and *Proteobacteria*

Unclassified *Bacteroidetes*, Unclassified *Cyclobacteriaceae*, Unclassified *Hymenobacteraceae*, and *Pontibacter* sp. are found under Bacteroidetes phylum. Except Unclassified Bacteroidetes, all others were found to have rich sets of carbohydrate active enzymes (Figure 3). Unclassified *Cyclobacteriaceae* showed presence of molecular signatures involved in reduction of nitrite to ammonia or nitric oxide. It also showed presence of pathway for release of nitrogen by denitrification of nitrous oxide. Bacteria belonging to *Hymenobacteraceae* showed presence of nitric oxide to nitrous oxide conversion and then release of nitrogen by denitrification of nitrous oxide. The results also reveal the phylum to be involved in sulfur metabolism by SOX complex.

In our study we found MAGs belonging to only two genera types of the Firmicutes phylum viz., Unclassified *Bacillaceae*, and *Exiguobacterium* sp. Both of them showed presence of PEGs to convert acetate to methane. They also hold few carbohydrates active enzymes reacting on polyphenolics, starch, chitin, pectin, galactans, and arabinan. They are rich in genes responsible for SCFA and alcohol conversions as shown in figure 3.

Proteobacteria is the most diverse group found on the test bacterial community as shown in figure 3, comprising of MAGs divided into 11 different clusters. Interestingly, majority of the organisms in this phylum have hydrocarbon metabolizing enzymes reacting on polyphenolics and chitin hence must have significant role in actual plastic degradation process. Except few unclassified alphaproteobacterial and one of unclassified *Rhodobacteraceae*, all other bacteria were holding potential to involve in nitrogen cycle. *Alteromonadaceae*, *Azoarcus pumilus*, *Gammaproteobacteria*, *Halomonas*, *Marinobacter*, *Rhodobacteraceae*, *Rhodobacterales*, *Thauera*, and *Zoogloeaceae* are also found to process sulfur using SOX complex. One of the *Rhodobacterales* is also having potential to reduce mercury. Except one gammaproteobacterial, all other bacteria are having methanogenesis or methanotrophy related genes. They also hold decent reaction of SCFA and alcohol conversions.

Based on the metabolic potential of the individual MAGs, we have functionally characterized the bacterial community and created a pattern as shown in figure 4a for representing potential interactions of bacterial community with the environment. We found that all phyla are involved in methanogenesis and nitrogen cycle. But only Bacteroidetes and Proteobacteria are involved in sulfur cycle and these genomes possesses genes involved in the SOX complex. According to the phylum distribution proteobacteria dominated the bacterial community. MAGs in the Proteobacteria taxa also possessed the arsenate and mercury reducing gene sequences. Proportion of Proteobacteria is lesser in control i.e., C as compared to PE-1 and PE-2 i.e., red and blue light treated respectively as shown in figure 4b. However, proportion of Bacteroidetes, Actinobacteria and Firmicutes in control is more as compared to light treated samples. Hence Proteobacteria must be a dominant part of plastic degrading bacterial community. Degradation of plastic is a tough task for bacteria. Therefore, bacteria surviving in extreme environments may be capable of handling such task. In this case, Proteobacteria MAGs are also predicted to possess heavy metals reduction ability like arsenate, mercury which is also a tough work. Plastics are of mainly two types which are either celluloid-based or phenolics-based (Jadhav et al., 2022). Phenolics-based plastics are even more thermotolerant as compared to celluloid-based plastics. A large number of Proteobacteria MAGs as observed showed potential presence of polyphenolics metabolizing pathways.

**Figure 4.**
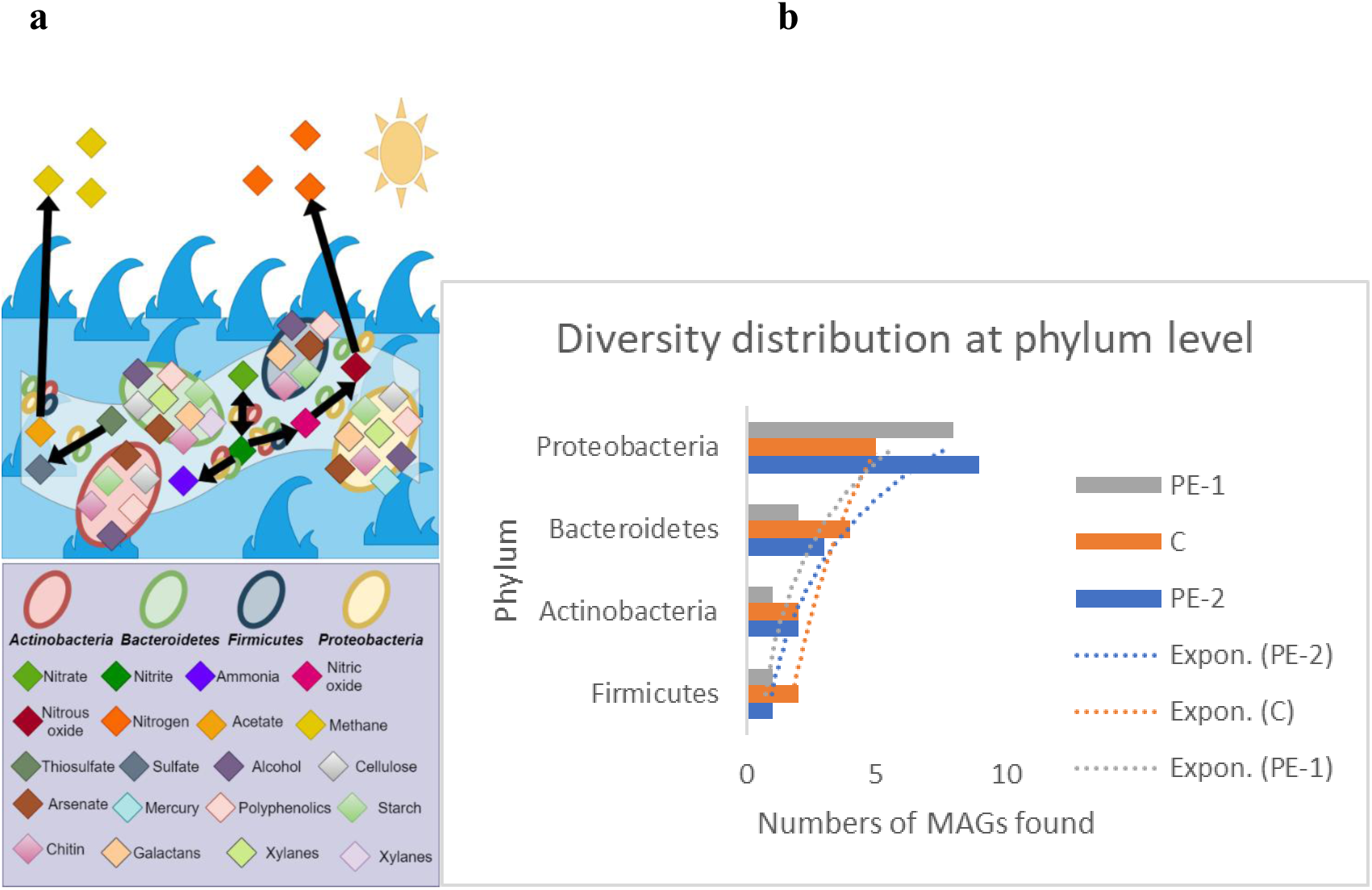
**a** Proposed model for plastic associated microbial community and their interaction with the surrounding environment, **b** Phylum level diversity distribution in the biofilm.

### Abundant bacteria found in plastic degrading bacterial community along with this community formation and inhibition mechanism

*Chromatiaceae* was found to detach abundantly and specifically in response to blue light (around 480nm) while *Piscirickettsiaceae* detach abundantly and specifically in response to red light (around 650 nm). These results indicating that photoreceptors in *Chromatiaceae* and *Piscirickettsiaceae* response to the blue and red-light wavelength electromagnetic radiations respectively. While *Cyclobacteriaceae*, *Rhodobacterales*, *Bacillaceae* separate abundantly from plastic associated bacteria in treated and untreated samples both. Hence, these three organisms not necessarily require treatment for detachment.

Statistical analysis was performed using STAMP software based on genome abundance. Abundantly obtained quality bins along with others subject to statistical analysis. Quality of the bins was accessed based on genome completeness (> 50%) and contamination (< 10%). Significant differences in bacteria with respective p-values in different samples. Horizontal bars in figure 5a indicated by blue, red and black colors indicating organisms detached from plastic by blue light, red light and without light (dark) treatment. Metagenomic alpha and beta diversity within and among samples are indicated in figure5 b-c.

**Figure 5:**
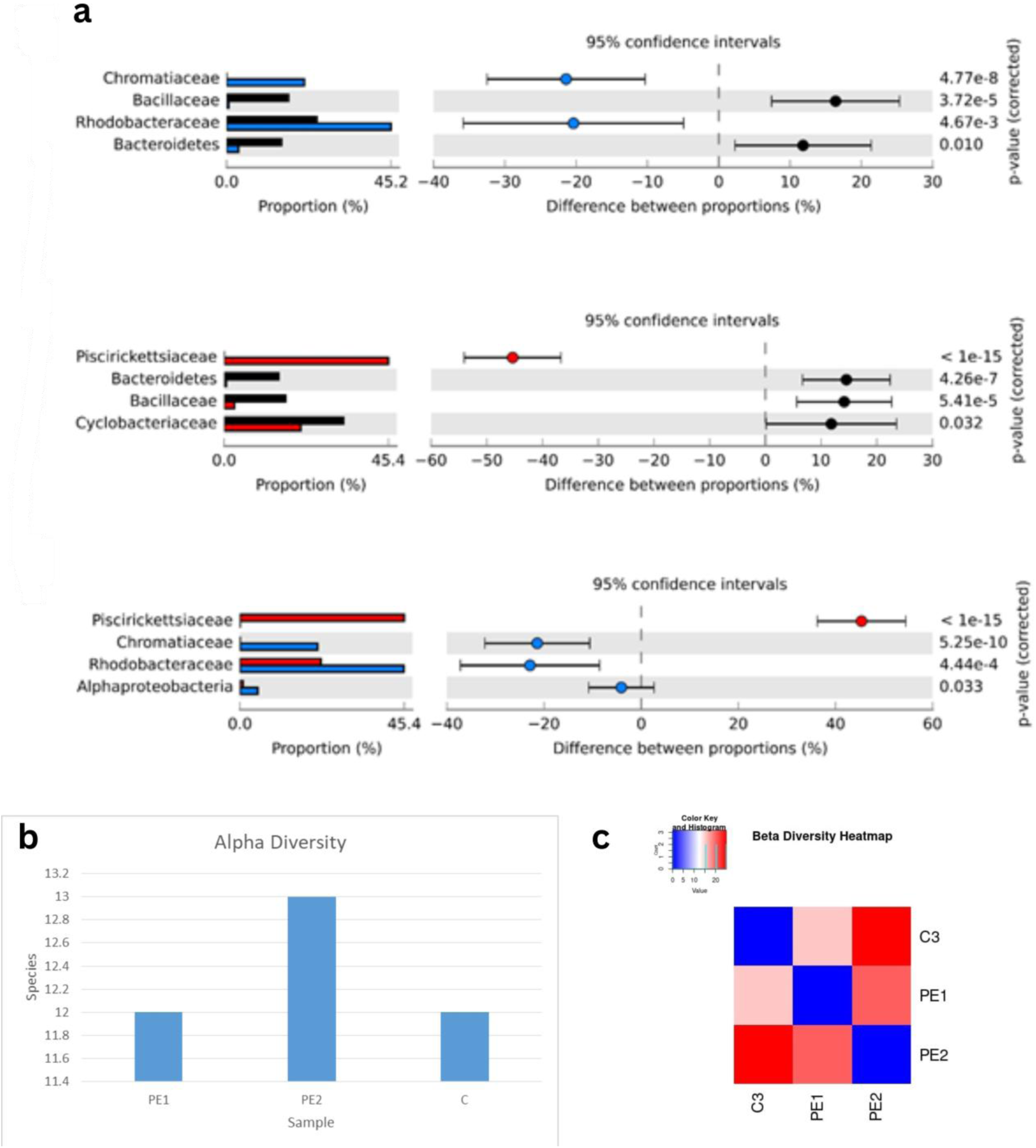
Metagenomic sample statistical analysis to check the differences among samples. a comparisons between blue light treated and control (without light treatment) samples; red light treated and control samples; red light treated and blue light treated samples respectively. b Alpha diversity within samples. c Beta diversity among samples.

### Limitations and future prospective

The only limitation of this study is while using electromagnetic radiations for bacterial community detachment might cause mutations in bacteria. Hence we might not get actual DNA sequence of the substrate attached bacterial community. This would affect research and development. However, this use of electromagnetic radiation strategy could be used for detachment of strongly attached pathogenic bacteria from their substrate. This would help in treatment of disease when using electromagnetic radiations in combinations of antibiotics. This technique would also be useful in study of plastic degrading enzymes and other genes involved in the degradation of plastics, since we can able to study exact bacterial species that can degrade the plastic.

## Conclusion

This study mainly deals with bacterial composition of plastic degrading bacterial community detached from plastic surface by electromagnetic radiation pretreatment (red and blue light only). Probable mechanism must include involvement of photoreceptor over bacterial surface or membrane which can able to sense the electromagnetic radiation of specific wavelength thereby activating gene inhibitor for plastic attachment bacterial genes and separate plastic degrading bacteria that response to particular light. Different specific bacteria were observed to be a part of plastic degrading bacterial community. These bacteria should involve in the process of plastic degradation, in association with other bacteria from plastic associated bacterial community. In this study, abundant plastic associated bacteria were *Cyclobacteriaceae*, *Rhodobacterales*, *Bacillaceae*, *Chromatiaceae* and *Piscirickettsiaceae*; in which *Piscirickettsiaceae* response specifically to red light and *Chromatiaceae* response specifically to blue light.

## Acknowledgement

Authors are grateful to the Director of CSIR-National Institute of Oceanography (CSIR-NIO), Goa, India and Scientist-in-Charge at CSIR-NIO, Regional Centre, Mumbai for their encouragement and support. HSJ is thankful to CSIR for the JRF fellowship.

## Ethical Approval

Not applicable.

## Consent to Participate

Not applicable.

## Consent to Publish

Not applicable.

## Authors Contributions

**ABF:** Conceptualization, Methodology, Writing- Reviewing and Editing, Funding Acquisition **HSJ:** Performed the sampling, Visualization, Formal analysis, Writing- Original draft preparation **CM:** Writing- Reviewing and Editing **NN:** Writing- Reviewing and Editing.

## Funding

This work was supported by OLP2009.

## Statements and Declarations

### Competing Interests

There is no competing interest.

### Availability of data and materials

Raw data is submitted to European Nucleotide Archive (ENA) under the study number PRJEB48191. MAGs are submitted to NCBI under bioproject ID PRJNA773851.

